# Nocturnal scent in a ‘bird-fig’: a cue to attract bats as additional dispersers?

**DOI:** 10.1101/418970

**Authors:** Simon P. Ripperger, Saskia Rehse, Stefanie Wacker, Elisabeth K. V. Kalko, Stefan Schulz, Bernal Rodriguez-Herrera, Manfred Ayasse

## Abstract

The plant genus *Ficus* is a keystone resource in tropical ecoystems. One of the unique features of figs is the diversity of fruit traits, which in many cases match their various dispersers, the so-called fruit syndromes. The classic example of this is the strong phenotypic differences found between figs with bat and bird dispersers (color, size, and presentation). The ‘bird-fig’ *Ficus colubrinae* represents an exception to this trend since it attracts the small frugivorous bat species *Ectophylla alba* at night, but during the day attracts bird visitors. Here we investigate the mechanism by which this ‘bird-fig’ attracts bats despite its fruit traits, which should appeal solely to birds. We performed feeding experiments with *Ectophylla alba* to assess the role of fruit scent in the detection of ripe fruits. *Ectophylla alba* was capable of finding ripe figs by scent alone under exclusion of other sensory cues. This suggests that scent is the main foraging cue for *Ectophylla alba.* Analyses of odor bouquets from the bat- and bird-dispersal phases (i.e. day and night) differed significantly in their composition of volatiles. The combination of these two findings raises the question whether *E. alba* and *F. colubrinae* resemble a co-adaptation that enables a phenotypically classic ‘bird-fig’ to attract bat dispersers by an olfactory signal at night thus maximizing dispersal.

---

Fruiting plants need to ensure that their seeds are transported away from their point of origin in order to increase survival probability by avoiding competition and reaching advantageous environments for germination [1]. Common ways of seed dispersal include self-dispersal by explosive fruits, dispersal by wind or the production of fleshy fruits to promote dispersal by animals [2]. Animal dispersal, or zoochory, frequently consists of a mutualistic relationship between plants and animals where animals are rewarded with edible, fleshy fruit parts for their service of transporting seeds away from the parental plant [3].

Bats and birds are very important vertebrate seed dispersers in tropical ecosystems [4, 5]. Fruits, however, that are consumed by either bats or birds may vary strongly in their appearance as a consequence of the contrasting sensory capacities and activity times of the associated dispersers [6]. Diurnal birds mainly rely on vision while foraging and hence prefer conspicuous fruits that contrast with the foliage [7-11]. On the contrary, bat fruits are frequently cryptic green and produce strong odors to attract their nocturnal dispersers [12-14]. Additionally, bat dispersed plants present fruits on erect spikes or pendulous structures in order to facilitate close distance detection by echolocation [14, 15]. While bats are able to consume larger fruits piecemeal by using their teeth, fruit size may be challenging to birds since they are limited by gape width [16-19].

Such different requirements of disperser groups have been suggested to drive the development of so-called dispersal syndromes, trait combinations that show a correlated evolution [20-22]. The existence of dispersal syndromes has been discussed for a long time and has recently received strong support by a recent study performed on Madagascar on lemur and bird dispersed fruits that showed an adaptation of scent production and composition only in lemur dispersed fruits [23] and by a comprehensive study of the plant genus *Ficus* [16, 24] a keystone resource for many tropical frugivores including bats and birds [12, 25]. In detail, bird dispersed figs or ‘bird-figs’ from both New and Old World tropics tend to be smaller, stronger contrasting to the foliage, less odorous, and arise from branches. On the contrary, figs dispersed mainly by bats or ‘bat-figs’ are larger, more cryptic relative to the foliage, have an aromatic scent, and are frequently presented on the trunk [6, 26].

The importance of olfaction for fruit detection in bats has been demonstrated in feeding trials for several frugivorous species of the Neotropical bat family Phyllostomidae [6, 13, 14, 26]. These studies show that the examined bat species are able to localize fruits by either olfaction alone or in combination with echolocation. This dominant role of olfaction in the foraging behavior of frugivorous bats may enable plants that phenotypically match the bird-dispersal syndrome to expand seed dispersal into the night by nocturnal production of volatiles that attract bats or other nocturnal mammals. The Paleotropical fig species, *Ficus benghalensis*, has been shown to produce significantly different odor bouquets during day and night, possibly in order to attract nocturnally foraging bats by scent, while diurnal birds are attracted by visual cues [27]. Unfortunately, the appeal of the altered scent on the nightly dispersers has not been studied in experimental setups.

The Mesoamerican fig species *F. colubrinae* is an excellent study organism to further investigate the mechanisms of attracting nightly dispersers despite heavy bird visits during day. The phenotype of *F. colubrinae* clearly matches the bird-dispersal syndrome with very small fruits which are bright red colored when ripe and presented on the branches [4, 28]. While birds extensively visit these fig trees during day, the small phyllostomid bat *E. alba* feeds heavily on fruits of *F. colubrinae* at night [29]. In the present study, we assess the role of fruit odor in the attraction of *E. alba* to ripe fruits and we characterize the diel variation in scent of fruits of *F. colubrinae*. If *F. colubrinae* would attract *E. alba* by nightly changes in scent, we would expect that the following hypotheses should be met: (1) olfaction plays a major role for the detection of ripe fruits in *E. alba*; (2) odor bouquets of fruits change when the fruits ripen and vary among day and night in ripe fruits, and (3) ripe fruits will shift production and release of volatiles during night in favor of substances that are known from published studies to be dominant in ‘bat-figs’. In order to test these hypotheses we combine semi-natural behavioral experiments with wild bats and chemical analyses of fig scent.

## METHODS

### Study site

Our study was conducted at „La Tirimbina Rainforest Center” (TRC) in the province Heredia in Costa Rica (10°26*’* N, 83°59*’* W). The study site is located in the Caribbean lowlands of Costa Rica. Annual precipitation averages at 3900 mm. Behavioral experiments were performed during May and June 2010 and sampling of fig scent from February to May 2011. Study organisms: *F. colubrinae* (Moraceae) is a Neotropical fig species. Its fruiting phenology is characterized by asynchronous fruit crop production of small fruits (diameter < 0.8 mm, mass 0.3 g) that are presented on the branches and turn dark red while ripening [12, 28]. On Barro Colorado Island in central Panama *F. colubrinae* draws little attention of frugivorous bats and is hence considered to be mainly bird-dispersed [12, 30]. However, farther north where *F. colubrinae* occurs in sympatry with *E. alba* this particular bat species shows a dietary specialization on *F. colubrinae* [29].

### Study animal

*E. alba* is a frugivorous, small-bodied leaf-nosed bat species (Phyllostomidae) that is distributed from northern Honduras to north-eastern Panama [31]. It modifies leaves, predominantly of plants of the genus *Heliconia*, to construct shelters where it roosts in social groups of typically four to eight individuals [29].

### Behavioral experiments

We captured groups of *E. alba* from roosts in *Heliconia* leaves in the area of TRC and selected single males for the feeding experiments in order to prevent lactating or pregnant females or juveniles from isolation of the social group. All individuals that were not considered for further experiments were set free immediately in close proximity to the roost. Following the capture, a single male was released into a flight tent (Eureka; ground area 4 × 4m, height 2.5m) several hours before sunset. At nightfall we installed a freshly cut branch of *F. colubrinae* that yielded a range of fruits of different stages of maturity into the flight tent. In order to adjust to the foraging situation we allowed the bat to feed on ripe fruits. After the consumption of five fruits we started choice trials in order to test whether *E. alba* relies mainly on olfaction for the short-range localization of ripe fruits or if objects similar in shape, color or presentation raise a bat’s attention. On one side of the branch we presented a strong olfactory cue to the bat that lacked visual or echo-acoustic properties of natural figs, i.e. we presented a fully opaque tissue bag that was filled with ten ripe figs (similar methods have been used to test for the response of bats to olfactory cues in absence of natural fruit shape or surface structure: Kalko and Condon (15) presented cotton saturated with juice of cucurbit fruits to bats; Hodgkison, Ayasse (26) wrapped ripe figs in several layers of nylon stockings). Simultaneously we presented on the other side of the branch odorless fig models made from red clay that were similar to natural *F. colubrinae* fruits in terms of form, color, and fruit presentation (in branch forks). We rated *E. alba*’s behavior as a positive response to the presented object when repeated approximation flights to or a landing next to the object followed by a directed movement to it occurred. In total, we tested six individual bats. Every bat was tested only once in order to avoid bias caused by learning effects. It was not possible to record data blind because our study involved focal animals. We documented bat behavior using an infrared camera (Sony Night-Shot DCR-HC42E, Sony, Japan) that was connected to a video recorder (GV-D 900E, Sony, Japan). We stored recordings on MiniDV video tapes (DVM60PR3, Sony, Japan).

### Sampling of fig scent

We sampled volatiles of *F. colubrinae* fruits based on dynamic headspace adsorption techniques [6, 26, 32]. Three categories of fruits were sampled: (1) unripe during night, (2) ripe during day, and (3) ripe during night. Fruits were collected from five individual fig trees and placed in glass chambers. Four glass chambers were connected to a single battery operated membrane pump. Every individual glass chamber was connected via a Teflon tube to an adsorbent tube containing activated charcoal (activated charcoal, Supelco, Orbo 32 large) that was installed upstream in order to filter-clean the pulled atmospheric air. After passing the glass chamber containing the fruit, the air exit through a glass sampling cartridge packed with 5mg Super Q (Waters Division of Millipore) in order to collect volatiles. The sampling cartridges were twice y-connected to the pump via silicone tubing. Two such setups were run simultaneously allowing for the collection of seven samples at a time along with one blank control that consisted of an empty glass chamber. Each sampling session was started at 2000 h for nightly sampling, or 0800 h for daily sampling, respectively, and lasted for eight hours with a flow rate of ca. 100mL min^−1^.

After sampling, all sorbent tubes were eluted with 0.050 ml of 10:1 pentane/acetone. Eluted samples were sealed in small airtight borosilicate glass specimen tubes and stored in the freezer at −18°C. After each sampling session, all glassware was thoroughly cleaned three times with ethanol (Absolute Alcohol, Hayman Ltd., Essex, UK), acetone (LiChrosolv, Merck, Darmstadt, Germany), and pentane (SupraSolv, Merck). Sorbent tubes were cleaned three times with ethanol, dichloromethane (LiChrosolv, Merck), and pentane, and then wrapped in aluminum foil and stored for future use in airtight glass jars with Teflon-coated lids.

### Chemical analyses of compounds

GC-Runs, Quantification & MS-Analyses: For quantitative analyses, 0.1 μg of octadecane was added as an internal standard to each of the eluted fruit odor samples collected by dynamic headspace adsorption (see above). All samples were analyzed with an HP5890 Series II gas chromatograph (Hewlett-Packard, Palo Alto, CA, USA), equipped with a DB5 capillary column (30 m × 0.25 mm i.d.) that used hydrogen as the carrier gas (2 ml min−1 constant flow). One microliter of each sample was injected splitless at 40°C. After 1 min, the split valve was opened and the temperature increased by 4°C min^−1^ until reaching a temperature of 300°C. GC/MS analyses were carried out on an HP 6890 Series GC connected to an HP 5973 mass selective detector (Hewlett-Packard) fitted with a BPX5 fused-silica column (25 m, 0.22 mm i.d., 0.25 μm film thick, SGE). Mass spectra (70 eV) were recorded in full scan mode. Retention indices were calculated from a homolog series of n-alkanes. Structural assignments were based on comparison of analytical data obtained with natural products and data reported in the literature [6, 26, 33], and those of synthetic reference compounds. Structures of identified candidate compounds were verified by co-injection.

### Statistical analyses

We performed principal component analysis (PCA) on the relative amounts of fruit scent compounds using SPSS 17. We used the resulting principal components (PCs) with an eigenvalue above one to run a discriminant function analysis (DFA) in order to test for differences in the scent composition between (1) unripe fruits during night, (2) ripe fruits during day, and (3) ripe fruits during night. We used the factor loadings after varimax rotation and the standardized discriminant function coefficients to assess the importance of individual compounds. Factor loading above 0.5 were considered high. Finally, we compared relative amounts of single compounds of ripe fruits during day and night (groups 2 and 3) using Mann-Whitney U-tests in R 2.15.3 [34].

## RESULTS

### Behavioral experiments

After releasing captured bats into the flight tent, the bats performed circular inspection flights for several minutes before they roosted in a corner of the flight tent until dusk. Shortly before dusk we installed a natural branch of *F. colubrinae* with several ripe and unripe fruits. All six bat individuals performed search flights that lasted between less than one minute and almost two hours (mean ± standard deviation: 32 ± 43 minutes, n = 6) until the bats approached the branch for the first time. Then the bats conducted two to nine approximation flight towards the branch over a period of one to 91 minutes (mean ± standard deviation: 19 ± 36 minutes, n = 6) before they landed and consumed a fig either directly on the branch or on the wall of the tent.

After the consumption of five ripe figs we started the behavioral experiments by presenting to the bat red modelling clay fig dummies on a natural branch of *F. colubrinae* and a tissue bag filled with 10 ripe *F. colubrinae* figs. None of the tested bats showed a clear positive response to the modelling clay figs. We did neither observe repeated approximation flights nor landing in the proximity of the models, which represented an echo-acoustic/visual cue similar to natural figs (a red, similar sized sphere presented in branch forks). On the contrary, five out of six individuals responded to the bag filled with ripe figs representing a strong olfactory cue (while one individual did not show any reaction to the experimental setup). After a period of six to 48 minutes (mean ± standard deviation: 16 ± 21 minutes, n = 5, see Table 1) and one to five approaches the bats either landed on or right next to the bag or landed more than 5 cm away and move hand over hand along the branch towards the bag. Subsequently the bats bit open the bag and consumed a fig.

**TABLE 1.**
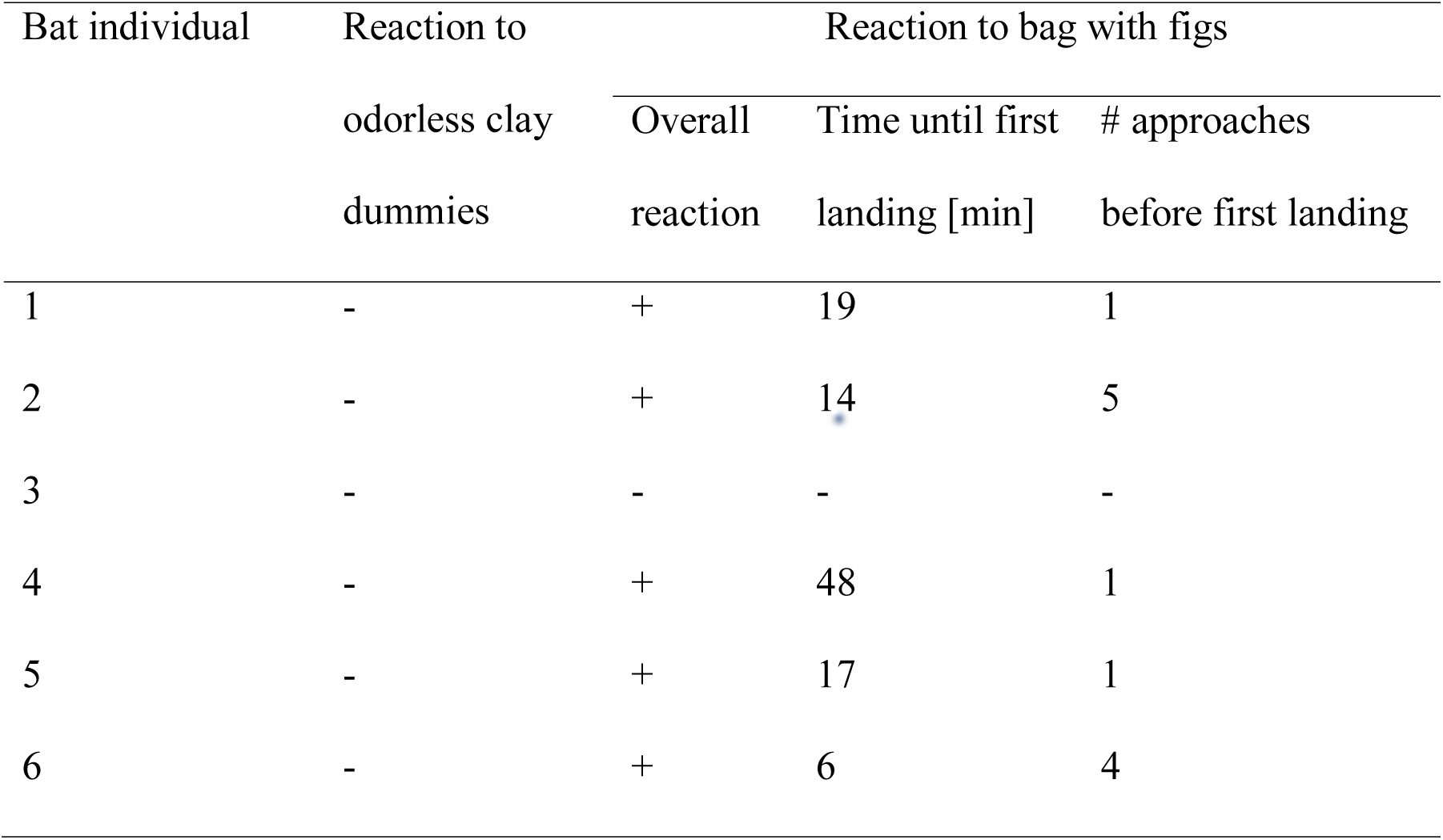
Parameters measured during behavioral trials on six individuals of *Ectophylla alba* that were subjected with odorless fig clay dummies and a bag filled with real scent releasing figs of *Ficus colubrinae*

### Chemical analyses

In the chemical analyses we registered 14 distinct peaks that were attributed to 17 individual substances (3 peaks showed co-eluting substances), 13 of which were unambiguously identified by mass spectrometry (Table 2). Nonanal and 1-tetradecanol contributed the largest share to the overall bouquet (Fig. 1, Table S1). Three further substances could be assigned to substance classes, however, not identified and one substance could not be classified. The identified substances belonged to different compound classes: aliphatic compounds derived from the fatty acid biosynthetic pathway (here shortly named fatty acid pathway compounds, FAPCs), sesquiterpenenes, and aromatic compounds. In three cases, two substances contributed to a single peak in the GC-analysis. In those cases, the overlapping substances were represented by a single value for the following analyses. Two of the identified substances, indene and anthracene, have a main relevance in industrial applications and were therefore excluded from all further analyses. They were considered environmental pollutants that accumulated on the outside of the fruits over time since our field site was closely located to human structures including infrastructure and industry. There were no significant differences in relative amounts of indene and anthracene among day and night in ripe fruits.

**FIGURE 1.**
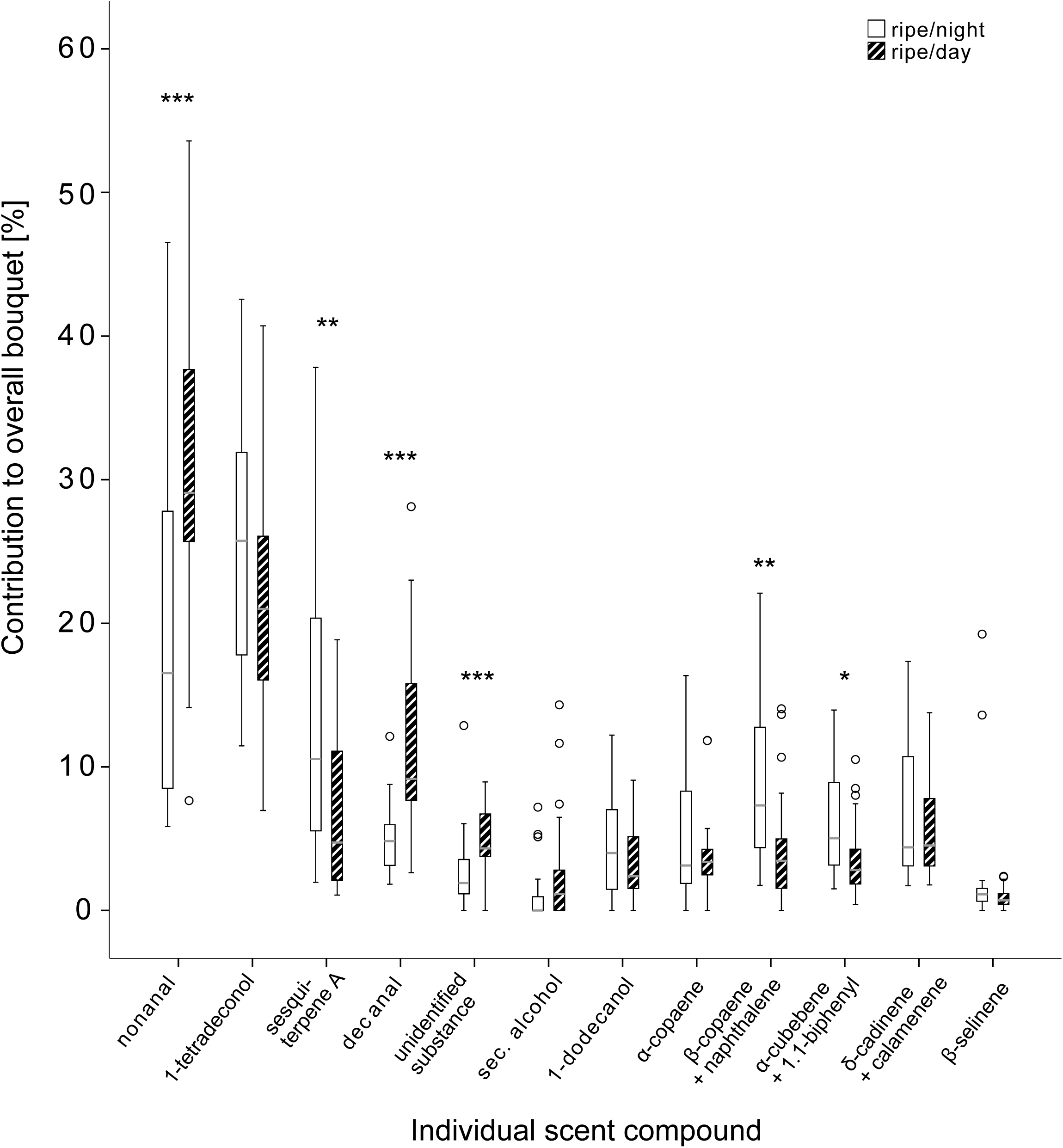
Relative amounts of compounds that contribute to the separation of ripe figs during daytime and night. Asterisks indicate significance based on the following α-levels: * p < 0.05, ** p < 0.01, *** p < 0.001

**TABLE 2.**
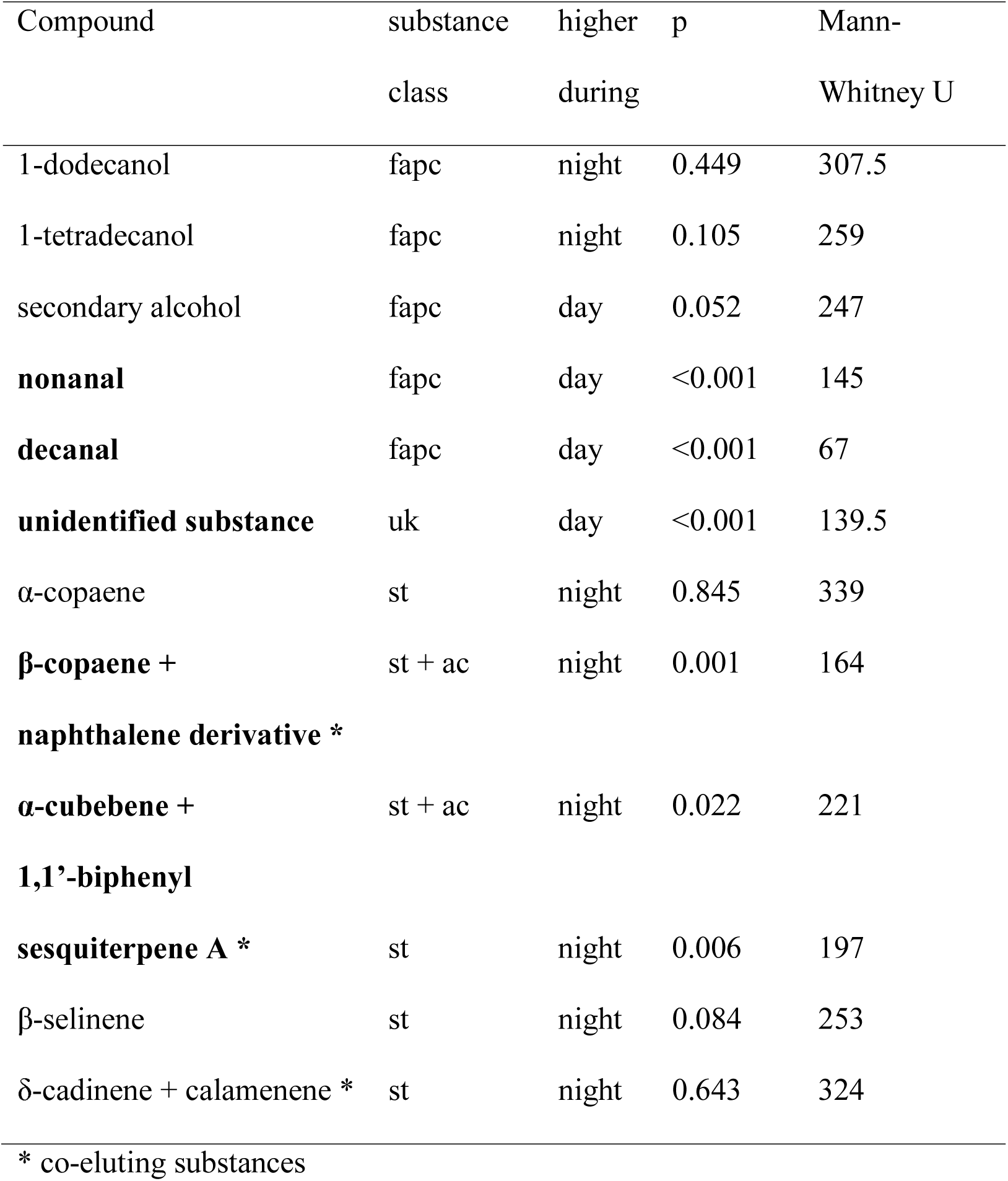
Comparison of individual chemical scent compounds of ripe fruits during day and during night based on relative amounts; fruit volatiles were attributed to the following classes: *fapc* fatty acid pathway compounds, *st* sesquiterpenes, *ac* aromatic compounds, *uk* unknown. Bold letters indicate significant differences between day and night (Mann-Whitney U).

We performed a PCA that included 12 individual values for the relative amounts of the remaining 15 chemical compounds from the three tested groups of figs ((1) unripe fruits at night, (2) ripe fruits during day, and (3) ripe fruits during night). Four PCs with an eigenvalue above one accounted for 76.2% of the total variation (see electronic supplement table S2). The DFA that used the four PCs as variables resulted in two discriminant functions (DFs, see table S3) and showed significant differences between the tested groups (function 1: χ² = 78.9, df = 8, p < 0.001; function 2: χ² = 24.9, df = 3, p < 0.001; Fig. 2). The highest coefficient for DF 1 was attributed to PC2, which in turn had high factor scores on the sesquiterpenes α-copaene and δ-cadinene + calamenene (Table S2 & Table S3). For DF 2, PC1 and PC3 had the highest coefficients. PC1 had high factor loading on sesquiterpene A, β-copaene + naphthalene derivative, α-cubebene + 1,1’-biphenyl and the FAPCs nonanal and decanal. 1-dodecanol and 1-tetradecanol loaded high on PC3. Seventy-five percent of the original grouped cases were correctly classified (72.5% of cross-validated grouped cases).

**FIGURE 2.**
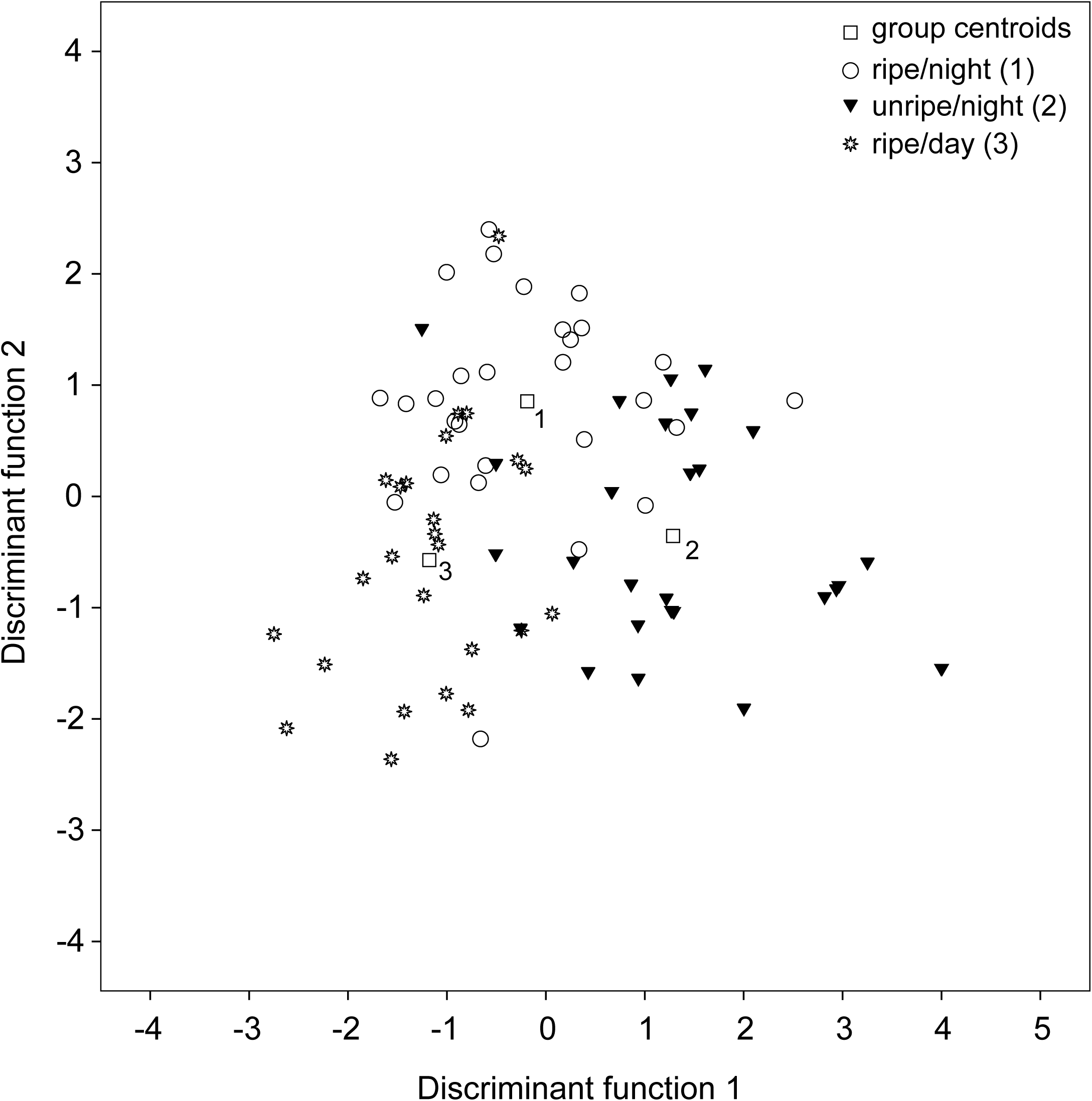
Comparison of scent bouquets produced by unripe fruits at night, ripe fruits at night and ripe fruits during day based on the composition of their chemical compounds using canonical discriminant function analysis (DFA)

Daily differences of single compounds in ripe fruits: All scent compounds analyzed were present in diurnal and nocturnal scents. In general, fatty acid pathway compounds dominated both diurnal and nocturnal scents (Fig. 1). However, relative amounts of sesquiterpene compounds increased at night and FAPCs decreased, except the two long-chain alcohols (Table 2). Six out of twelve day/night comparisons of relative amounts of single scent components showed significant differences. The aldehydes nonanal and decanal and one unclassified substance accounted for a significant greater share during day, while three sesquiterpene compounds in combination with aromatic compounds (sesquiterpene A, β-copaene + naphthalene derivative, α-cubebene + 1.1-biphenyl) had significantly higher proportions during night (Table 2).

## DISCUSSION

Our study shows that scent seems to be an important cue for *E. alba* to find ripe fruits of *F. colubrinae*. *Ectophylla alba* was capable during experimental trials to find ripe figs by scent alone under exclusion of other natural sensory cues. Odor bouquets of figs undergo significant changes with regard to the relative amounts of compounds during the process of maturation, and in our chemical analyses we found that bouquets of ripe figs differ significantly in the composition of volatiles during day and night. Nightly changes in scent composition show a pattern that contrasts with other ‘bat-figs’, but some compound of the ‘night-bouquet’ have been reported in other fruits consumed by small phyllostomid bats. We suggest that our findings show initial signs of an adaptation of *F. colubrinae* towards dispersal by small bats such as *E. alba*, but further behavioral trials are required to unequivocally demonstrate this relationship.

Semi-natural feeding trials showed that phyllostomid bats locate fruits by echolocation [15] or olfaction [6, 13, 14] as the primary sensory cues. Our results from the feeding experiments suggest that *E. alba* conforms to the latter foraging strategy. The tested bats only showed strong responses to the tissue bag that gave a strong olfactory cue but lacked natural texture, shape, size, or presentation of figs that might be of importance for detection by echolocation. Therefore, we assume that echolocation may not play such a dominant role for *E. alba* in fruit detection as it does for other bat species. The Neotropical bat *Phyllostomus hastatus* feeds on fruits of a Cucurbitaceae that are borne on pendulous structures [15]. This style of fruit presentation facilitates detection by echolocating bats because the fruit represents a clutter free target. In general, flagellichory or cauliflory (pendulous or trunk-borne presentation of fruits that reduce the presence of foliage close to the fruit) are widespread adaptations of plants to chiropterochory [35]. Korine and Kalko (13) argue that detection of fruits by downwards frequency modulated signals which are typical for phyllostomid bats is possible but largely depends on the fruit presentation and the complexity of the surrounding clutter. However, *F. colubrinae* presents its fruits sessile, usually paired at the node [28], thus in a highly cluttered environment making detection by echolocation difficult. Hence, *F. colubrinae*’s way of fruit presentation may favor olfaction as primary cue for detecting figs, at least until *E. alba* gets very close to the figs.

Olfactory cues enable plants to signal the readiness of fruits for dispersal. Accordingly, temporal changes in the volatile profile of fruits are common during the process of ripening (e.g. [36-38]) and have also been documented for wild, bat-dispersed fig species [26]. Our data are consistent with a change in the overall composition of the scent bouquet during the process of ripening. Additionally, we observed significant changes among day and night, caused by day-time specific scent production. Circadian changes in the volatile profile of fruits have been more rarely observed than changes during ripening. To our knowledge, only Borges, Ranganathan (27) observed diel differences in the volatile signal in Old World figs of the species *F. benghalensis*. These fruits are consumed by birds during the day and by bats during the night. Dispersal by both, birds and bats, is not uncommon within the genus *Ficus*, yet this dispersal mode usually concurs with fruit phenotypes that are considered intermediate between the bird and the bat syndrome [24]. While most fruit traits in *F. colubrinae* match the bird-syndrome, scent alone can be sufficient for *E. alba* to detect the ripe fruits as shown by our behavioral experiments. This observation raises the question whether a nightly shift in volatile production may enable ‘bird-figs’ to additionally attract certain bat species as dispersers and hence allow for dispersal during the daytime and at nighttime. To achieve seed dispersal by distinct animal taxa may result in multiple benefits to a reproducing plant. The contribution to overall seed rain by birds or bats, respectively, may vary quantitatively across seasons [4]. Microhabitat deposition also strongly depends on the disperser since birds tend to disseminate seeds when perched while bats usually defecate seeds during flight. The resulting seed rain can be dominated by chiropterochorously dispersed seeds at forest edges and open areas, while most ornithochorous seeds reach forest sites [39, 40]. An all-season reproducing plant species like *F. colubrinae* that may develop both, epiphytic and solitary life forms [28], may in particular benefit from the attraction of both bats and birds. This way the plant may maximize dispersal rates of the year-round produced fruits and seeds may arrive in a more heterogeneous range of microhabitats for germination.

All unambiguously identified compounds except 1-dodecanol, 1-tetradecanol, and calamenene have been documented to be produced by *Ficus* spp., either by floral stages (Grison-Pigé, Hossaert-McKey (41): α-cubebene, α-, β-copaene, β-selinene, δ-cadinene, decanal) or by fruits (Hodgkison, Ayasse (6): α-, β-copaene, δ-cadinene; Borges, Ranganathan (27): nonanal, decanal, α-copaene, δ-cadinene). The scent bouquet of *F. colubrinae* fruits, which was dominated by fatty acid pathway compounds, was more similar to ‘bat-figs’ from the Old World tropics [26, 27, 42] than to Neotropical bat-dispersed fig species that were characterized by high proportions of monoterpenes [6]. Monoterpenes were completely missing in our samples. This result was surprising since feeding trials showed that fruit scents, which were dominated by monoterpenes were highly attractive to the phyllostomid bat *Artibeus jamaicensis* [6]. Instead, in our samples sesquiterpenes increased throughout and in parts significantly during night, while fruit scents that were dominated by sesquiterpenes were rejected by *A. jamaicensis*. Interestingly, sesquiterpenes, including α- and β-copaene, dominated the bouquet of the only small sized Neotropical fig species (*F. costaricana*) in the sample of Hodgkison [6], a ‘bird-fig’ the seeds of which can occasionally be found in the feces of small bat species [30, 43]. Calamenene, α-copaene and β selinene have further been identified from the scent of inflorescences of *Calyptrogyne ghiesbreghtiana* [44]. This palm is visited by bats including small *Artibeus* species (*watsoni/phaeotis*) [45], which also feed on small-sized figs [30]. These differences observed among figs and other bat-dispersed plants raise the question whether different olfactory preferences exist in bats that have different diets, as it was already proposed by Hodgkison and colleagues [6].

## CONCLUSIONS

Taking the results from behavioral trials and chemical analyses together, our study cannot finally answer the question, whether a ‘bird-fig’ like *F. colubrinae* attracts additional, nightly dispersers by altered scent production, but it provides evidence that this might indeed be a strategy in the genus *Ficus*. Daily variation in the volatile profile of fruits may be more common than previously thought, since it has now been documented in both the New and the Old World tropics and volatile ecology in the genus *Ficus* seems to be complex and worth to receive further attention. It remains unclear whether daily variation in scent profiles is simply a consequence of plant physiology or a co-adaption among plants and dispersers. Future behavioral experiments that present bats with diurnal versus nocturnal scent bouquets of fruits might help to answer the aforementioned question, ideally including multiple species of bats and figs.

## ACKNOWLEDGMENTS

We thank Manuel Rojas, Christian Schmid, and Wito Lapinski for their help during field work. We are grateful to Gabriele Wiest for quantitative analysis of gas chromatograms. Emma Berdan, Thomas Blankers, and Linus Günther helped to improve this manuscript with well-conceived comments. Necessary permits were obtained with the valuable assistance of J. Guevara. The project was funded by the Deutsche Forschungsgemeinschaft (DFG, Germany) to EKVK and MA (KA 124 8-1; AY 12/8-2).

## DATA AVAILABILITY STATEMENT

Data will by archived upon article acceptance.

