## Supplement tables 1-3 for "Nocturnal scent in a ‘bird-fig’: a cue to attract bats as additional dispersers?"

**Table S1** Median and mean absolute deviation (mad) of relative amounts of 12 chemical compounds. Scent was collected from unripe fruits during night and from ripe fruits during night or day

| | unripe during night | | ripe during night | | ripe during day | |
|---|---|---|---|---|---|---|
| Substance | median | mad | median | mad | median | mad |
| 1-dodecanol | 4.7 | 2.71 | 4.01 | 4.11 | 2.4 | 1.6 |
| 1-tetradecanol | 15.18 | 3.9 | 25.75 | 11.19 | 21 | 7.52 |
| secondary alcohol | 0 | 0 | 0 | 0 | 1.16 | 1.72 |
| Nonanal | 18.83 | 10.04 | 16.54 | 12.27 | 29.07 | 9.62 |
| Decanal | 6.14 | 4.67 | 4.84 | 2.16 | 9.17 | 4.94 |
| unidentified substance | 2.64 | 1.35 | 1.92 | 2.03 | 4.33 | 2.64 |
| α-copaene | 10.88 | 8.87 | 3.14 | 2.63 | 3.38 | 1.33 |
| β-copaene + naphtalene derivative | 7.41 | 4.7 | 7.32 | 5.09 | 3.46 | 2.83 |
| α-cubebene + 1,1’-biphenyl | 2.73 | 2.48 | 5.04 | 3.74 | 2.82 | 1.89 |
| sesquiterpene A | 6.84 | 4.42 | 10.56 | 9.06 | 4.74 | 4.51 |
| β-selinene | 2.4 | 1.33 | 1.13 | 0.65 | 0.71 | 0.58 |
| δ-cadinene + calamenene | 12.25 | 7.65 | 4.4 | 2.18 | 4.53 | 2.54 |

**Table S2** Factor loadings of 12 volatile compounds on four principal components; factor loadings > 0.5 are shown

| | PC1 | PC2 | PC3 | PC4 |
|---|---|---|---|---|
| β-copaene + naphthalene derivative | 0.913 | | | |
| sesquiterpene A | 0.912 | | | |
| α-cubebene + 1,1’-biphenyl | 0.857 | | | |
| nonanal | -0.733 | | | |
| decanal | -0.727 | | | |
| α-copaene | | 0.825 | | |
| δ-cadinene + calamenene | | 0.789 | | |
| 1-tetradecanol | | | -0.819 | |
| 1-dodecanol | | | -0.656 | |
| secondary alcohol | | | | 0.836 |
| unidentified substance | | | | 0.662 |
| Cumulative proportion of variance | 31.5 | 50.3 | 63.6 | 76.2 |

**Table S3** Standardized canonical discriminant function coefficients of four principal components

| principal component | function | |
|---|---|---|
| | 1 | 2 |
| 1 | 0.366 | -0.737 |
| 2 | 0.948 | 0.293 |
| 3 | 0.059 | 0.663 |
| 4 | -0.562 | 0.298 |
